# Cell Cycle Control by Nuclear Sequestration of *CDC20* and *CDH1* mRNA in Plant Stem Cells

**DOI:** 10.1101/198978

**Authors:** Weibing Yang, Raymond Wightman, Elliot M. Meyerowitz

## Abstract

In eukaryotic cells, most RNA molecules are exported into the cytoplasm after being transcribed in the nucleus. Long noncoding RNAs (lncRNAs) have been found to reside and function primarily inside the nucleus, but nuclear localization of protein-coding messenger RNAs (mRNAs) has been considered rare in both animals and plants. Here we show that two mRNAs, transcribed from the *CDC20* and *CCS52B* (plant orthologue of *CDH1*) genes, are specifically sequestered inside the nucleus during the cell cycle. CDC20 and CDH1 both function as coactivators of the anaphase-promoting complex or cyclosome (APC/C) E3 ligase to trigger cyclin B (C YCB) destruction. In the *Arabidopsis thaliana* shoot apical meristem (SAM), we find *CDC20* and *CCS52B* are co-expressed with *CYCBs* in mitotic cells. *CYCB* transcripts can be exported and translated, whereas *CDC20* and *CCS52B* mRNAs are strictly confined to the nucleus at prophase and the cognate proteins are not translated until the redistribution of the mRNAs to the cytoplasm after nuclear envelope breakdown (NEBD) at prometaphase. The 5’ untranslated region (UTR) is necessary and sufficient for *CDC20* mRNA nuclear localization as well as protein translation. Mitotic enrichment of *CDC20* and *CCS52B* transcripts enables the timely and rapid activation of APC/C, while their nuclear sequestration at prophase appears to protect cyclins from precocious degradation.

## Introduction

Understanding the patterns and regulatory mechanisms of organ formation in multicellular organisms is a central aspect of developmental biology (Lander, 2011). Animal organogenesis is completed during embryonic development or, in some instances, during metamorphosis; while in plants, active division and differentiation of stem cells and their progenitors in the shoot apical meristem (SAM) and the root apical meristem (RAM) lead to continuous formation of new tissues and organs, ensuring developmental plasticity in a changing environment (Gaillochet and Lohmann, 2015; Heidstra and Sabatini, 2014; Meyerowitz, 1997; Vernoux et al., 2000). Plant cell division, as in mammalian cells, yeast and *Drosophila*, is triggered and maintained by the kinase complex composed of cyclin-dependent kinases (CDKs) and various cyclin subunits. Fluctuating gene expression and orderly proteolysis of cyclins, spatial positive feedback of Cdk1-cyclin B1 redistribution, combined with the antagonistic actions of Wee1 kinase and Cdc25 phosphatase, generate a robust and highly ordered mitotic process (Coudreuse and Nurse, 2010; Dewitte and Murray, 2003; De Veylder et al., 2007; Morgan, 1995; Santos et al., 2012).

Destruction of cyclins at the appropriate time in the cell cycle is mediated by APC/C, an E3 ubiquitin ligase whose catalytic activity and substrate specificity are conferred by two coactivators, CDC20 and Cdc20 homolog 1 (CDH1) (Peters, 2006; Pines, 2011; Yu, 2007). During early mitosis, phosphorylation of the APC/C subunits, such as the auto-inhibitory segment loop in APC1, exposes the binding sites of CDC20 thus facilitating CDC20 association with APC/C (Fujimitsu et al., 2016; Kraft et al., 2003; Qiao et al., 2016; Zhang et al., 2016). At prometaphase, APC/C activity is restrained by the spindle assembly checkpoint (SAC), a regulatory pathway during which unattached kinetochores generate a diffusible ‘wait anaphase’ signal that triggers the incorporation of CDC20 into a complex composed of MAD2, BUBR1 and BUB3, leading to the formation of the mitotic checkpoint complex (MCC) (Fraschini et al., 2001; Hardwick et al., 2000; Sudakin et al., 2001). Recently it has been proposed that MCC itself could function as a diffusible signal to inhibit APC/C by recognizing a second CDC20 that has already bound to and activated APC/C (Izawa and Pines, 2015). Furthermore, APC/C activity is counteracted by the F box protein early mitotic inhibitor 1 (Emi1) (Reimann et al., 2001). The multi-faceted regulation of APC/C in various organisms suggests high plasticity of APC/C activity, and also implies the existence of additional mechanisms.

Subcellular RNA localization has been implicated in multiple cellular processes by regulation of spatial gene expression (Lipshitz and Smibert, 2000). For instance, the posterior-anterior polar localization of *bicoid*, *oskar*, *gurken*, and *nanos* mRNAs in *Drosophila* oocytes guides proper pattern formation and embryo development (Martin and Ephrussi, 2009). Long noncoding RNAs (lncRNAs) predominantly localize to the nucleus to modulate transcription factor binding, histone modification, chromosome structures and specific nuclear body formation (Batista and Chang, 2013; Engreitz et al, 2016; Geisler and Coller, 2013; Tsai et al., 2010). While mature mRNAs are considered to reside predominantly in the cytoplasm, deep sequencing of nuclear and cytoplasmic RNA fractions from various mouse tissues identified a number of mRNAs with higher amounts in the nucleus than in the cytoplasm (Bahar Halpern et al., 2015), suggesting a potential for mRNA nuclear retention in gene expression regulation. However, nuclear localization of mRNAs or mRNA precursors and its biological relevance have rarely been documented. In *Drosophila* embryos, the non-polyadenylated *histone* mRNAs are retained in the nuclei of DNA-damaged cells, contributing to the maintenance of genome integrity (Iampietro et al., 2014). *CTN-RNA*, an adenosine-to-inosine (A-to-I) edited mouse-specific pre-mRNA, localizes in the nuclear paraspeckle and can be rapidly cleaved under physiologic stress to produce *mCAT2* mRNA encoding a cell-surface L-arginine receptor (Prasanth et al, 2005). Apart from these examples, nuclear sequestration of non-edited mature mRNAs remains to be discovered.

Here, through a comprehensive fluorescent in situ hybridisation (FISH) analysis of mRNA distribution of core cell cycle genes in *Arabidopsis* stem cells, we have found that *CDC20* and *CDH1* orthologue *CCS52B* mRNAs are sequestered in the nucleus during prophase. We show that *CDC20* and *CCS52B* transcripts accumulate to peak levels but are confined to the nucleus at prophase, and redistribute into the cytoplasm following NEBD at prometaphase. With fluorescence live cell imaging, we demonstrate that this mRNA nuclear sequestration blocks CDC20 and CCS52B protein translation, thus preventing premature APC/C activation in early mitosis. By systematic mRNA deletion and chimeric RNA localization analysis, we found that *CDC20* mRNA 5’UTR confers nuclear sequestration and is also involved in protein translation. Nuclear sequestration of *CDC20* and *CCS52B* mRNAs reveals a previously unrecognized mechanism for the tuning of APC/C activity.

## Results

### Systematic Analysis of mRNA Localization of Core Cell Cycle Genes in Meristematic Cells

In *Arabidopsis*, the SAM is organized into three zones distinguished by cell division activity: the central zone (CZ) composed of slowly dividing stem cells, which is surrounded by the peripheral zone (PZ) that contains rapidly dividing cells that give rise to primordia of leaves and flowers, and the rib meristem (RM) underlying the CZ and the PZ responsible for stem growth (Steeves and Sussex, 1989) (Figure1A). The distinct cell division activities in different SAM regions can be visualized by using a fusion of green fluorescent protein (GFP) to CyclinB1;1 (CYCB1;1-GFP), exhibiting a low number of GFP-positive cells in the slowly dividing cells of the CZ and RM, and relatively higher number in the PZ and flower primordia (Figure 1B). Using a GFP-microtubule-binding domain marker (GFP-MBD) and the nuclear reporter histone H2B fused to red fluorescent protein (H2B-RFP), we found that the microtubule and nuclear structures corresponding to different cell cycle stages could all be identified in the SAM (Figure 1C). Therefore, the SAM serves as a suitable system with which to study the control of the cell cycle in plants.

**Figure 1.**
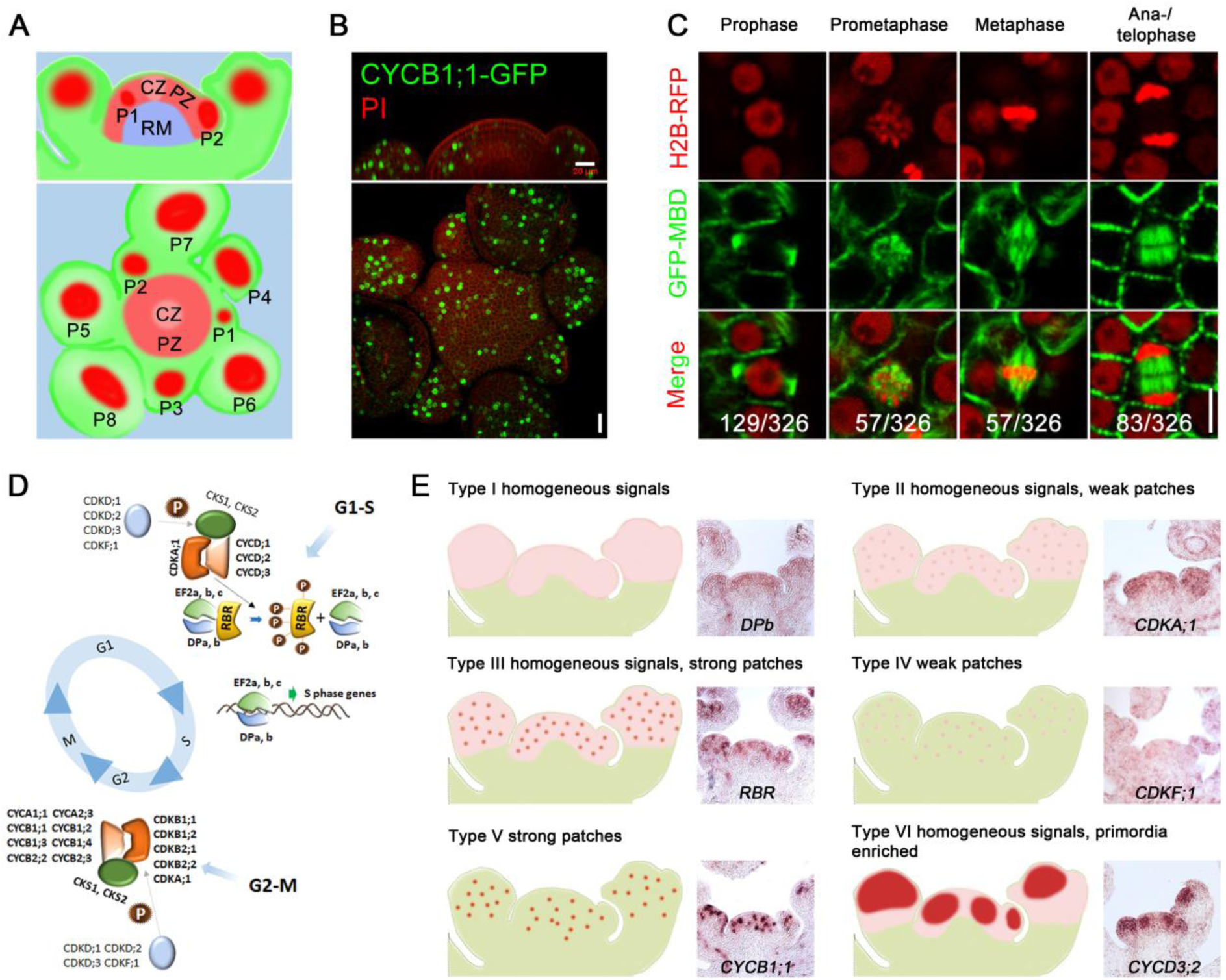
Expression Patterns of Core Cell Cycle Genes in *Arabidopsis* Meristematic Cells. (A) A schematic representation showing the organization of the *Arabidopsis* inflorescence shoot apical meristem (SAM). Upper panel, side view; lower panel, top view. CZ, central zone; PZ, peripheral zone; RM, rib meristem; P, flower primordia, which form sequentially in the PZ. (B) CYCB1;1-GFP reporter expression in wild type (WT) SAM. Scale bar, 20 µm. (C) Expression of nuclear reporter H2B-RFP and microtubule reporter GFP-MBD in meristematic cells corresponding to different cell cycle stages. From 6 WT SAMs, 326 cells were observed to be undergoing division and the number of cells at each stage is shown. Scale bar, 5 µm. (D) Functional modules of core cell cycle regulators in the *Arabidopsis* SAM. (E) Classification of the mRNA distribution patterns of core cell cycle genes expressed in the SAM. *In situ* hybridisation images for representative genes in each class are shown.

CDKs, CYCs and other regulatory proteins constitute a group of core cell-cycle regulators. Multiple members in each CDK and cyclin subfamily exist in plants, suggesting a level of functional conservation but also specialized regulation of cell cycle progression in plants as compared to animals (Vandepoele et al., 2002) (Figure 1D). To explore the role of cell cycle regulatory genes in *Arabidopsis* SAM development, we first analysed their mRNA abundance from RNA-seq data of meristematic cells derived from dissection of enlarged *clavata3* (*clv3*) mutant SAM (Yang et al., 2016). We focused on 130 annotated core cell-cycle regulators (Menges et al., 2005; Van Leene et al., 2010), and identified 72 genes showing detectable expression in the SAM (TPM > 10; Table S1). To investigate their expression pattern *in planta*, we carried out systematic RNA *in situ* hybridization. Using RNA probes specific to individual SAM-expressed cell cycle genes, we were able to detect the distribution of transcripts from 66 genes at single-cell resolution. In situ hybridization results are presented in Data S1. Most of the genes exhibited strong expression in the SAM compared to other tissues (e.g. stem), supporting the RNA-seq data. Based upon their expression patterns, these cell-cycle genes were classified into six groups: (i) homogeneous signal (Type I); (ii) homogeneous background signal with weak additional signal in a spotted pattern (Type II); (iii) homogeneous background signal with strong additional spots of signal (Type III); (iv) weak spots of signal in a subset of cells (Type IV); (v) only strong spots of signal in a subset of cells (Type V); and (vi) homogeneous background expression with additional strong signal in developing primordia (Type VI) (Figure 1E; Table S1). Homogeneous signals across the whole meristem indicate that the corresponding genes are expressed throughout the cell cycle; whereas patchy patterns suggest that expression correlates to specific cell cycle stages.

Most of the G1/S regulators, including *CDKA;1*, *E2Fs* (*E2Fa*, *E2Fb* and *E2Fc*) and *DPs* (*DPa* and *DPb*), displayed homogeneous expression in the shoot apex, which would maintain these meristematic cells with the capacity for active proliferation. One exception was *RETINOBLASTOMA RELATED* (*RBR*), an inhibitor of E2F and DP transcription factors, which showed a strong patchy pattern (Type III) (Figure 1E), similar to previous observations in embryonic and root meristematic cells (Wildwater et al., 2005), and implying a cell-cycle controlled regulation. Compared to G1/S genes, G2/M regulators, including plant-specific B type CDKs (*CDKBs*), and A and B type cyclins (*CYCAs* and *CYCBs*) were all grouped into Type V, showing a strongly patchy pattern with weak background expression (Figure 1E). RNA fluorescence *in situ* hybridization (RNA FISH) together with 4’, 6-diamidino-2-phenylindole (DAPI) staining indicated that these genes were exclusively expressed in mitotic cells from early prophase until late anaphase (Figure S1).

Our gene expression map data were consistent with Affymetrix microarray data of dividing *Arabidopsis* cell cultures (Menges et al., 2005). The mRNA distribution patterns in the shoot apex, combined with previous cell-cycle transcript *in situ* analysis in *Arabidopsis* seedlings and in the shoot/floral meristems of *Antirrhinum majus* (de Almeida Engler et al., 2009; Fobert et al., 1994), provide a good overview of cell cycle gene expression patterns in various plant tissues.

### Mitosis-specific Expression of *CDC20* and *CCS52B* mRNA

The accumulation of *CYCB* transcripts at M-phase (Figures S2A and S2B) would be expected to lead to a corresponding peak of CYCB proteins at this stage. Indeed, CYCB1;1-GFP fluorescence signals increased from prophase onwards, peaked at metaphase and then decreasing rapidly at anaphase, finally being undetectable in telophase cells (Figures S2C-S2E). The decline of CYCB1;1-GFP fluorescence signals could be caused by insufficient protein synthesis and/or short half-life. The rapid elimination of large amount of CYCB1 proteins may attribute to APC/C-mediated degradation (Figure 2A), a mechanism conserved among various organisms. The genes encoding *Arabidopsis* APC/C subunits, as well as the *CDH1* orthologues *CCS52A1* and *CCS52A2*, were all expressed homogeneously in the SAM at relatively low level (Table S1 and Data S1). By contrast, both *CDC20* and *CCS52B* showed strong patchy patterns similar to *CYCB* genes (Data S1). The distinct expression patterns of A- and B-class *CCS52* genes supported the predicted roles of CCS52As in regulating endoreduplication (Cebolla et al., 1999; Lammens et al., 2008; Vanstraelen et al., 2009), and *CCS52B* in controlling mitosis (Tarayre et al., 2004).

**Figure 2.**
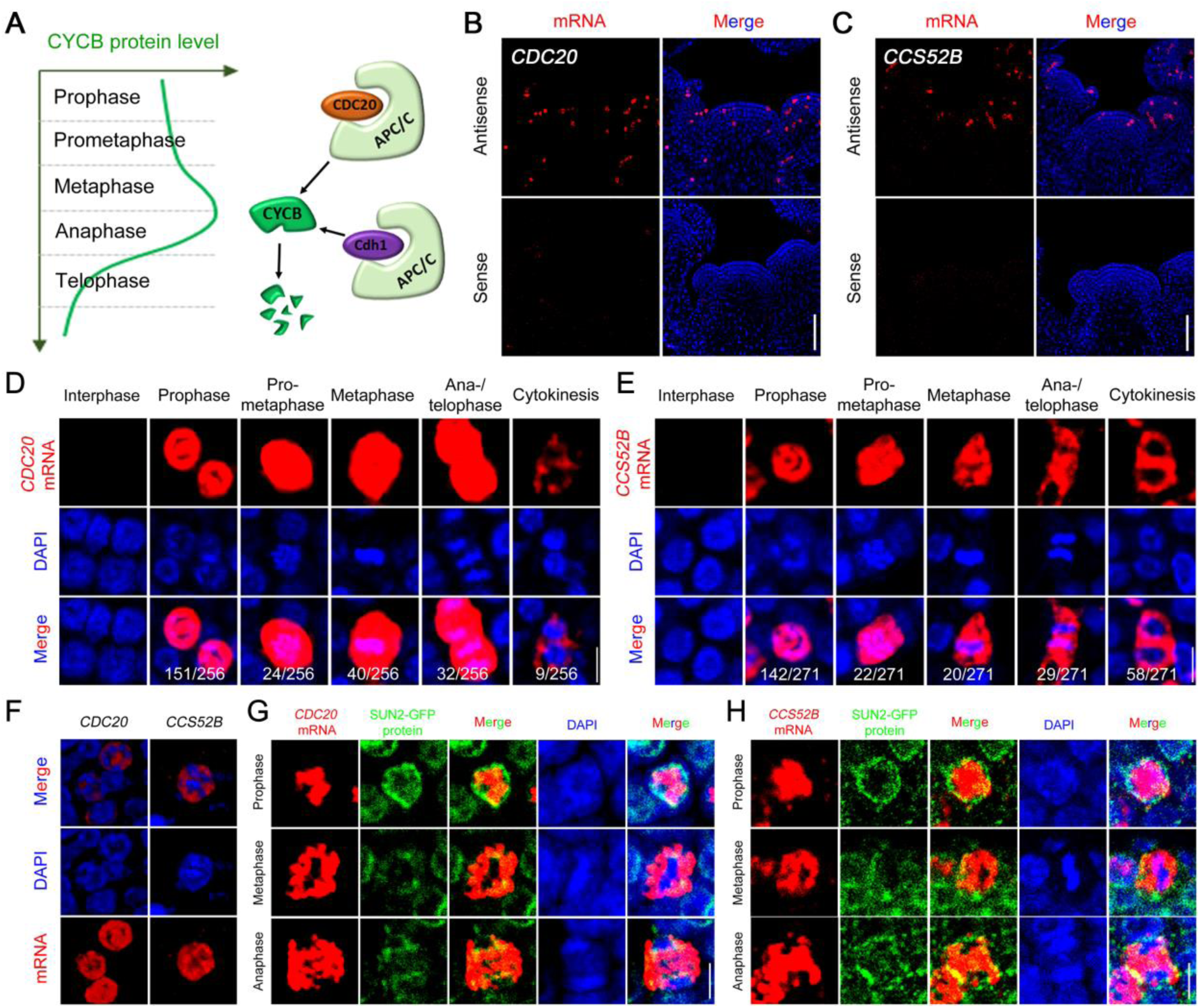
Nuclear Sequestration of *CDC20* and *CCS52B* mRNAs in Prophase Cells. (A) A schematic model illustrating CYCB protein dynamics during mitosis and its degradation by APC/C^CDC20^ and APC/C^CDH^1^^ E3 ligases. (B and C) RNA FISH to reveal the expression patterns of *CDC20* and *CCS52B* in the SAM. No signals were detected from the sense probes. Scale bars, 50 µm. (D and E) Expression of *CDC20* and *CCS52B* at different mitotic stages. Note the nuclear localization of *CDC20* and *CCS52B* mRNAs at prophase. Scale bars, 5 µm. (F) 3-D projection of *CDC20* and *CCS52B* mRNAs in prophase cells. Scale bar, 5 µm. (G and H) *CDC20* and *CCS52B* mRNA localization with nuclear envelope reporter at different stages of mitosis. The mRNAs were detected by FISH. The nuclear envelope was revealed using GFP antibody against a nuclear envelope reporter protein, SUN2-GFP. Scale bars, 5 µm. All images, with the exception of (F), show single optical confocal sections.

Cell-cycle controlled *CDC20* and *CCS52B* expression was further investigated by RNA FISH. Both mRNAs accumulated exclusively in mitotic cells from prophase until cytokinesis (Figures 2B-2E). The amount of *CDC20* mRNA decreased when mitosis was completed (Figure 2D), whereas a high level of *CCS52B* mRNA persisted until cytokinesis (Figure 2E). The extended expression of *CCS52B* relative to *CDC20* was validated by double RNA FISH. *CDC20* and *CCS52B* mRNAs co-expressed in early mitotic cells, but at late mitosis a population of cells were found only to express *CCS52B* (Figure S3). Taken together, the enrichment of *CDC20* and *CCS52B* transcripts, along with the constitutive expression of all APC/C components, would presumably allow for rapid APC/C activation.

### *CDC20* and *CCS52B* mRNAs Are Sequestered in the Nucleus at Prophase

Mature mRNAs are usually rapidly exported out of the nucleus (Köhler and Hurt, 2007). For example, *CYCB* transcripts, despite their high levels, were all found to reside in the cytoplasmic space (Figures S1 and S2). However, when analysing the sub-cellular distribution of *CDC20* and *CCS52B* mRNAs in prophase cells, we found that each of them is localized inside the DAPI-labelled nuclei (Figure 2F). No hybridization signals could be detected in the cytoplasm even when we increased the detection settings to saturation (data not shown). To further validate the nuclear sequestration of *CDC20* and *CCS52B* transcripts, we examined *CDC20* and *CCS52B* mRNA localization in mitotic cells together with a marker for the nuclear envelope. *CDC20* and *CCS52B* mRNAs were detected by RNA FISH. The nuclear envelope was revealed by immunohistochemistry using an anti-GFP antibody in SAM sections of *Arabidopsis* nuclear envelope marker line SUN2-GFP (Oda and Fukuda, 2011; Varas et al, 2015). As shown in Figures 2G and 2H, both *CDC20* and *CCS52B* mRNAs were localized inside the nucleus and were surrounded by the intact nuclear envelope in prophase cells; when cells enter metaphase and the nuclear envelope has disassembled, the transcripts were found distributed in the cytoplasm. At late telophase and cytokinesis when the nuclear envelope reforms, *CDC20* and *CCS52B* mRNAs were excluded from the nuclei of daughter cells, suggesting that once in the cytoplasm, *CDC20* and *CCS52B* mRNAs are not imported back or recruited into the nucleus. These cytosol-localized *CDC20* and *CCS52B* mRNAs seem to be unstable as they could only be detected in a small group of newly divided cells. Nuclear localization of *CDC20* mRNA was also detected in root apical meristem (Figures S4A and S4B) and shoot vascular cambium (Figure S4C), demonstrating that this phenomenon exists in the dividing cells of different tissues.

### Nucleo-cytoplasmic Compartmentalization of *CDC20* and *CYCB* mRNAs

Since both *CYCBs* and *CDC20* transcripts could be detected at prophase, we hypothesized that they might be expressed simultaneously in the same cells, although the possibility of sequential expression could not be excluded. To clarify this, we investigated *CYCBs* and *CDC20* expression in the same meristems by double RNA FISH. *Arabidopsis* wild-type meristems were hybridised with both *CYCBs* and *CDC20* gene-specific RNA probes and the number of cells expressing different genes was quantified. *CDC20* was found to largely co-express with different *CYCB* genes in all mitotic cells from prophase until anaphase (Figures 3A and 3C), whereas no co-expression was detected for *CDC20* with the S phase marker *Histone H4* (*HIS4*) gene (Figure 3B). In prophase cells, the localization of *CDC20* and *CYCB* transcripts was clearly separated: *CDC20* mRNA was restricted to the nuclei and surrounded by cytoplasmically localized *CYCB* mRNAs (Figures 3A and S5). Therefore, *CYCB* mRNAs can be translated, resulting in high expression of CYCB1;1-GFP in prophase cells (Figure S2); whereas nuclear confinement of *CDC20* and *CCS52B* transcripts might prevent protein synthesis.

**Figure 3.**
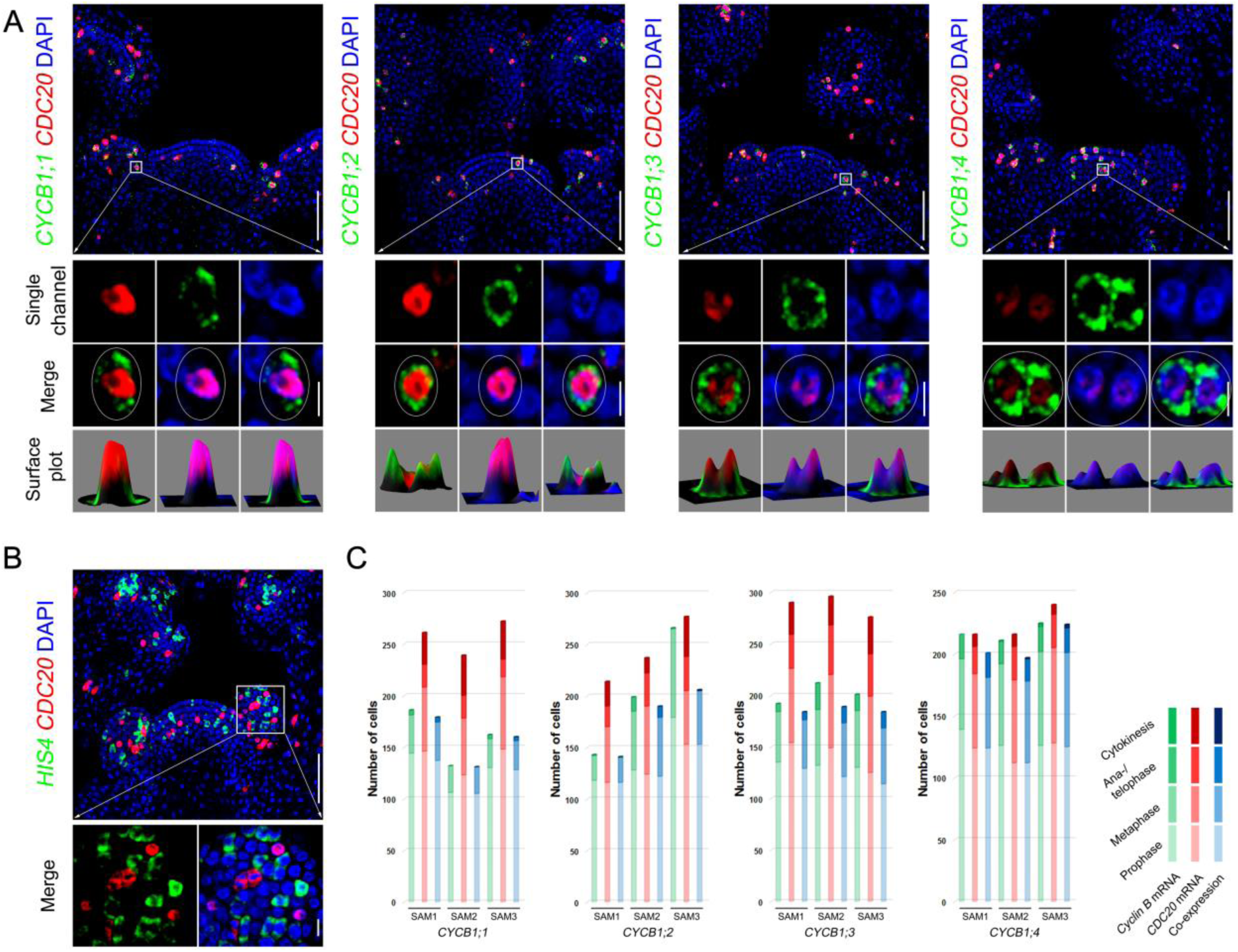
Spatial Separation of *CDC20* and *CYCB* mRNAs in Prophase Cells. (A) Co-expression of *CDC20* with cell cycle genes as revealed by double RNA FISH coupled with DAPI staining. (B) *CDC20* does not co-express with an S-phase expressed gene *HIS4*. CDC20 and cell cycle genes were detected by gene specific probes with different labelling. Scale bars in (A) and (B), SAM overview (top panels) = 50 µm; single cells (bottom panels) = 5 µm. (C) Quantification of the number of cells that express *CDC20* and *CYCB* genes at different mitotic stages. *CYCB1* genes were mostly expressed at prophase and metaphase, and largely co-express with CDC20.

### Nuclear Sequestration of *CDC20* and *CCS52B* mRNAs Blocks Protein Translation

To evaluate the effect of *CDC20* and *CCS52B* mRNA nuclear sequestration upon protein translation, we analysed the expression patterns of GFP-tagged CDC20 and CCS52B fusion proteins in living cells, an approach that has been widely used to track the dynamics of key cell cycle proteins, including CDC20 in animal cells (Nilsson et al., 2008). Genomic fragments containing the entire coding sequences of *CDC20* and *CCS52B* were fused with *GFP* at the N-terminus and expressed in wild-type plants under the control of their endogenous promoters. Double RNA FISH using *GFP* probe and *CDC20* and *CCS52B* gene-specific probes showed overlapping signals at different mitotic stages, suggesting that fusion of *GFP* coding sequence did not interfere with nuclear localization of *CDC20* or *CCS52B* mRNAs (Figure S6).

The meristems of p*CDC20::GFP-*g*CDC20* and p*CCS52B::GFP-*g*CCS52B* transgenic plants were examined using confocal microscopy. GFP-CDC20 was only expressed in a small fraction of meristematic cells (Figure 4A). GFP-CCS52B protein expression could be identified in a greater proportion of SAM cells, which predominantly localized in the nucleus but also in the cytoplasm (Figure 4C). For both proteins, the expression levels varied between different cells (Figures 4B and 4D). To analyse their expression in relation to different phases of the cell cycle, we further introduced GFP-CDC20 and GFP-CCS52B into H2B-RFP plants. GFPCDC20 fluorescence signals were detected at very low level in prometaphase cells, increased slowly at metaphase and anaphase, and reached maximal level in late telophase cells. When cytokinesis was completed, GFP-CDC20 eventually decreased and disappeared (Figures 4E and 4F). Compared to GFP-CDC20, the expression of GFP-CCS52B was much delayed, as it was not detected until cells enter late telophase. GFP-CCS52B protein expression exhibited its peak level at cytokinesis, and persisted until the next G1 stage (Figures 4G and 4H).

**Figure 4.**
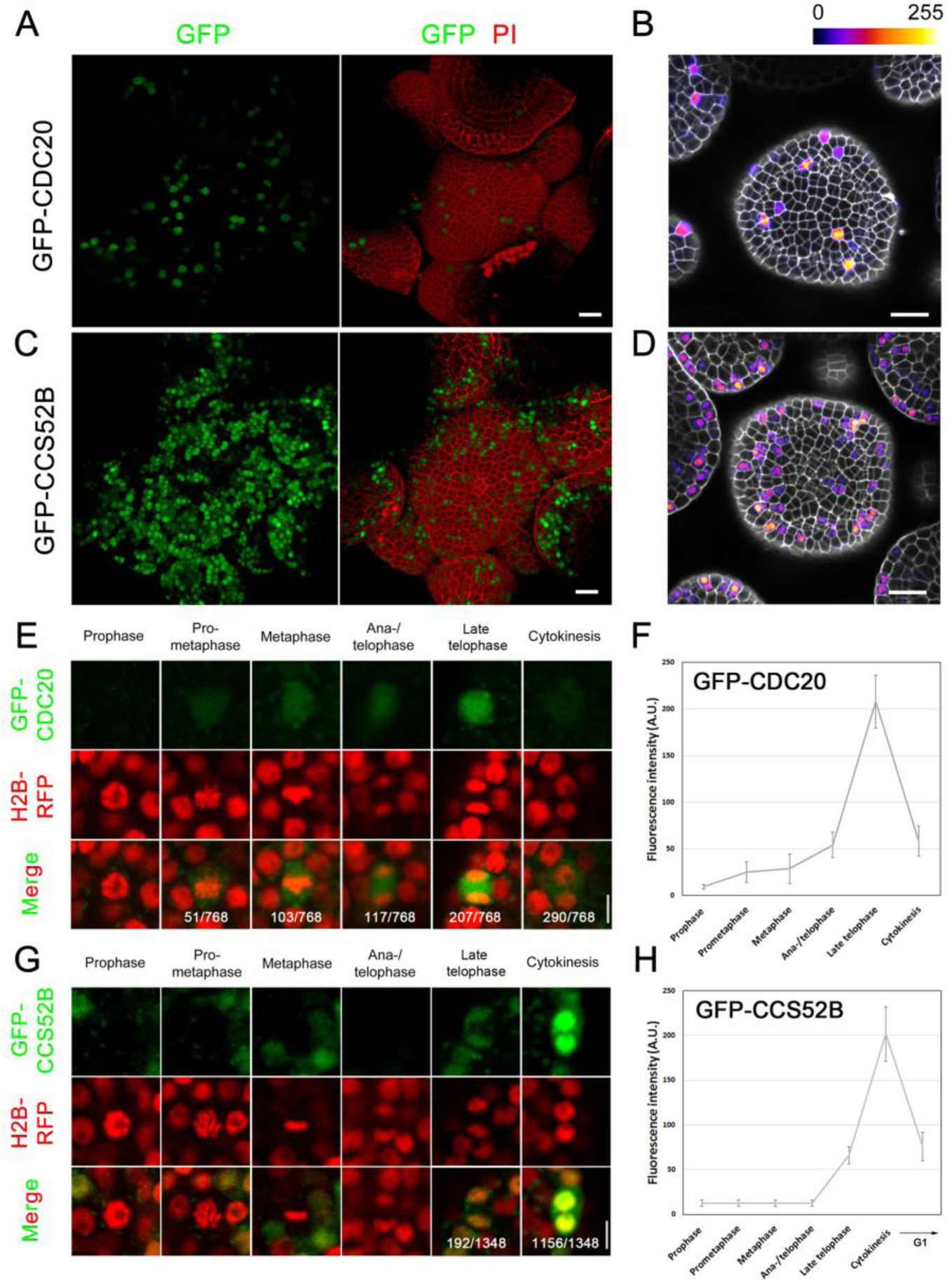
Expression Patterns of CDC20 and CCS52B Proteins during the Cell Cycle. (A-D) GFP-CDC20 (A, B) and GFP-CCS52B (C, D) expression in the *Arabidopsis* SAM. The cell wall was stained with propidium iodide (PI). Expression of GFP-CDC20 and GFPCCS52B in (B) and (D) were displayed using the Fire lookup table in ImageJ to show difference in fluorescence intensity. Scale bars, 20 µm. (E-H) Protein dynamics of GFP-CDC20 (E) and GFP-CCS52B (G) at different stages of mitosis. The fluorescence intensity was shown in (F) and (H). Scale bars, 5 µm.

The protein expression pattern of CDC20 beginning at prometaphase was consistent with its transcript accumulation prior to NEBD, followed by mRNA redistribution into the cytoplasm after NEBD. However, given the late appearance of CCS52B protein despite much earlier release of its RNA from the nucleus, it appears that there are additional mechanisms beyond nuclear sequestration that controls CCS52B translation, one of which could be regulation by *CCS52B* mRNA binding proteins as RNA-binding proteins also play crucial roles in controlling translation efficiency besides guiding RNA localization (Lipshitz and Smibert, 2000). Nevertheless, the peak accumulation of CCS52B protein at cytokinesis and subsequent stages was in line with the predicted roles of Cdh1 to degrade CDC20 and maintain a low cyclin abundance through late mitosis and G1 phases (Fang et al., 1998). After analysing a number of meristems from different transgenic lines, we were unable to detect any GFP-CDC20 or GFPCCS52B protein expression in prophase cells, demonstrating that mRNA nuclear sequestration correlated with an absence of protein translation.

### Dynamic Turnover of CDC20 and CCS52B proteins

The GFP-CDC20 and GFP-CCS52B proteins dynamics was further examined by real-time fluorescence imaging of individual cells, revealing that both proteins accumulated rapidly at late mitosis and disappeared when mitosis was completed (Figures S7A and S7B). Fluctuation in CDC20 protein levels during the cell cycle has been observed in animal cells (Fang et al., 1998; Kramer et al., 1998; Prinz et al. 1998). For CDH1, the protein level appears to remain constant throughout the cell cycle in HeLa cells (Fang et al., 1998; Kramer et al., 1998). In order to distinguish changes in gene expression from proteolytic activity, we treated SAMs with the proteasome inhibitor MG132. This treatment did not increase the protein level of GFPCCS52B, suggesting that CCS52B levels are a function of gene expression and translation (Figure S7C). By contrast, MG132 treatment resulted in a marked increase in GFP-CDC20 fluorescence intensity in both SAM and root cells (Figures S7D and S7E), implying that CDC20 may undergo continuous synthesis and degradation. Therefore, a conserved surveillance system exists to tightly control CDC20 protein abundance in plants as in human cells (Ge et al., 2009; Izawa and Pines, 2015; Nilsson et al., 2008).

### **Mapping the Cis-acting Element Involved in *CDC20* mRNA Nuclear Localization**

To investigate how *CDC20* mRNA is sequestered in the nucleus, we first tested the mechanisms proposed for known nuclear RNAs. It has been shown that mRNAs containing adenosine (A)-to-inosine (I) edited *Alu* inverted repeats are predominantly localized in the nucleus (Chen et al., 2008). A-to-I editing was responsible for the nuclear retention of *CTNRNA* (Prasanth et al., 2005). We compared the sequences of *CDC20* full-length cDNA and genomic DNA but did not find any difference, ruling out A-to-I editing in *CDC2*0 mRNA. In addition, mRNA transcribed from *CDC20* cDNA, like the genomic DNA-derived mRNA, was also confined to the nucleus (Figure S8A), suggesting that *CDC20* nuclear sequestration can act upon the mature mRNA. These results indicate that the regulation of *CDC20* mRNA nuclear localization was distinct from other nuclear RNAs.

As the targeting signals of localized RNAs are usually encoded by their own sequences (Buxbaum et al., 2015), we next sought to identify the cis-acting element involved in *CDC20* mRNA nuclear localization. A series of deletions spanning the entire *CDC20* coding sequence, each 200 bp in length (except for Δ1207-1374) with 100 bp overlapping, were fused with *GFP* and expressed in wild-type plants under the control of the *CDC20* promoter (Figure 5A). The localization of these truncated *GFP-CDC20* chimeric RNAs was examined by RNA FISH. As cytoplasmic localization of *CDC20* mRNA can be observed at late mitosis when daughter cell nuclei reform (Figure 2D), we used *CYCB1;2* mRNA expression as an indicator of prophase cells. *CYCB1;2* showed similar expression in these transgenic plants compared to wild-type plants (Figure 5B), suggesting that expression of these exogenous RNAs did not interfere with normal cell cycle progression. Examination of the subcellular distribution revealed that all these *GFP-CDC20* truncated RNAs were all localized inside the nucleus, surrounded by the cytoplasmic *CYCB1;2* mRNA (Figures 5B and S8B), indicating that deletion of a single fragment of *CDC20* coding region was not sufficient to disrupt RNA nuclear localization.

**Figure 5.**
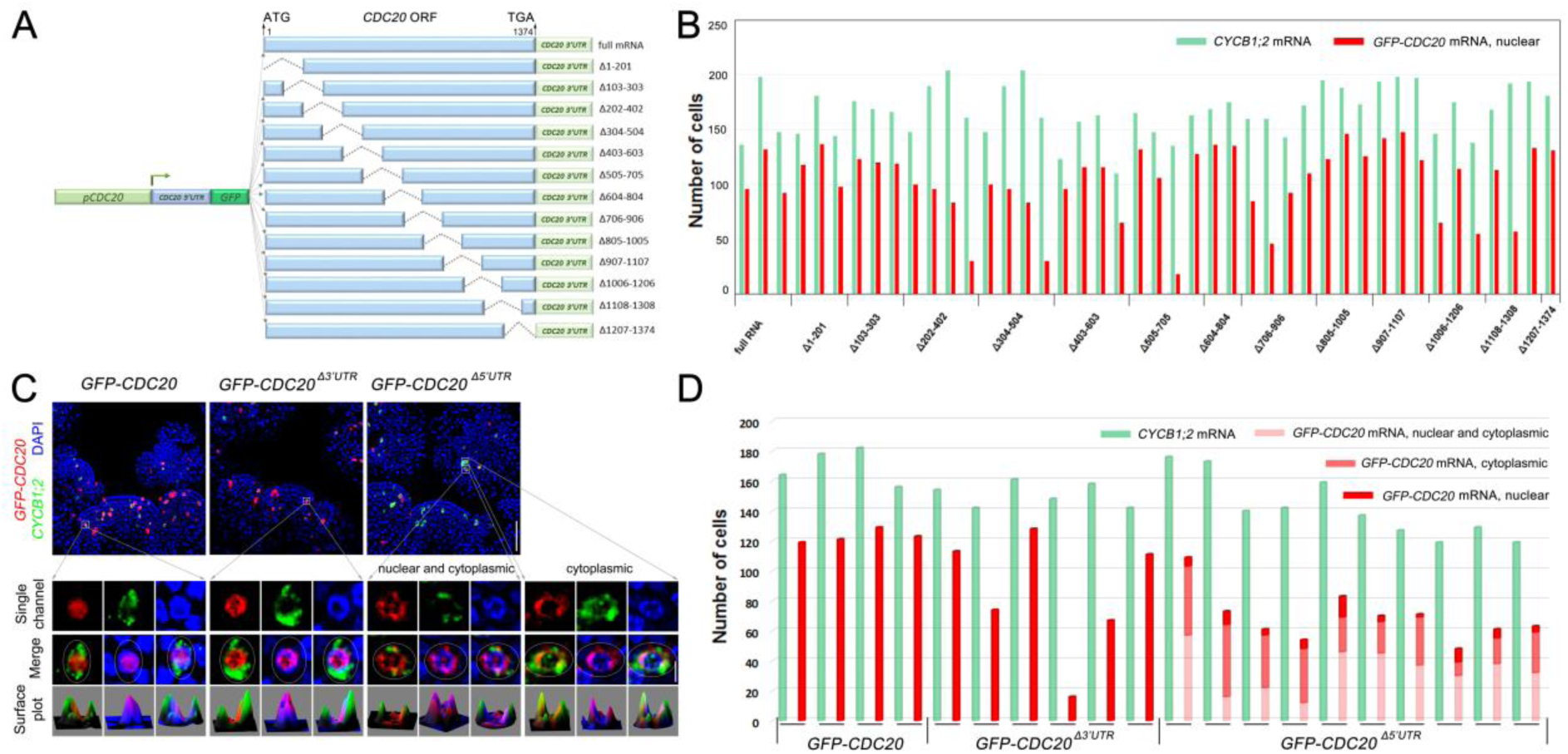
*CDC20* 5’UTR Is Involved in mRNA Nuclear Localization. (A) Schematic diagram of *CDC20* mRNA deletion constructs. (B) Quantification of the number of prophase cells expressing *GFP* fused *CDC20* mRNAs that contain serial deletions. *CYCB1;2* expression was used as a prophase marker. All *GFP-CDC20* mRNAs with deletions in the *CDC20* ORF were found to localize in the nucleus. Each pair of columns represents data from one meristem. (C) Localization of *GFP-CDC20* truncated mRNAs lacking *CDC20* 5’UTR or 3’UTR. Deletion of 5’UTR abolished *GFP-CDC20* mRNA nuclear sequestration, leading to nucleocytoplasmic or mostly cytoplasmic localization. Scale bars, 50 µm for SAM overview (top panels) and 5 µm for single cells (bottom panels). (D) Quantification of the number of prophase cells expressing full length, 3’UTR deleted, and 5’UTR deleted *GFP*-*CDC20* mRNAs. Each pair of columns represents data from one meristem.

We next investigated the role of UTRs in *CDC20* mRNA nuclear sequestration. Chimeric mRNAs transcribed from *GFP* in-frame fused with *CDC20* genomic fragment without the 5’UTR or 3’UTR (*pCDC20::GFP-CDC20^Δ5’UTR^* and *pCDC20::GFP-CDC20^Δ3’UTR^*) were analysed by RNA FISH. *CDC20* 3’UTR-truncated mRNAs showed the same nuclear localization pattern as full length *GFP-CDC20* transcript. By contrast, when the 5’UTR was deleted, nuclear localization was largely reduced. In most of the prophase cells, 5’UTR-truncated *GFP-CDC20* mRNAs were present either in both the nucleus and the cytoplasm, or mostly in the cytoplasm (Figures 5C, 5D and S8B), indicating that deletion of the 5’UTR abolished *CDC20* mRNA nuclear sequestration.

### *CDC20* 5’UTR Is Sufficient to Confer Nuclear Sequestration

To further evaluate the function of the *CDC20* 5’UTR, we fused it to a *GFP* coding sequence (Figure 6A). This chimeric mRNA, *5’UTR^CDC20^-GFP*, as well as *GFP* alone, were expressed in wild-type plants under the control of the *CDC20* promoter. The number of prophase cells expressing these *GFP* mRNAs seem to be reduced compared to *GFP* fused with full length *CDC20* mRNA (Figure 6C), implying that the *CDC20* coding region contains cis-element contributing to transcriptional activity. Nevertheless, when expressed, *5’UTR^CDC20^-GFP* mRNA was found to be exclusively confined to the nucleus. The control, *GFP* mRNA alone, was distributed in the cytoplasm similar to *CYCB1;2* mRNA (Figures 6B and S8C). The results, taken together, demonstrate that the 5’UTR was both necessary and sufficient to sequester *CDC20* mRNA inside the nucleus.

**Figure 6.**
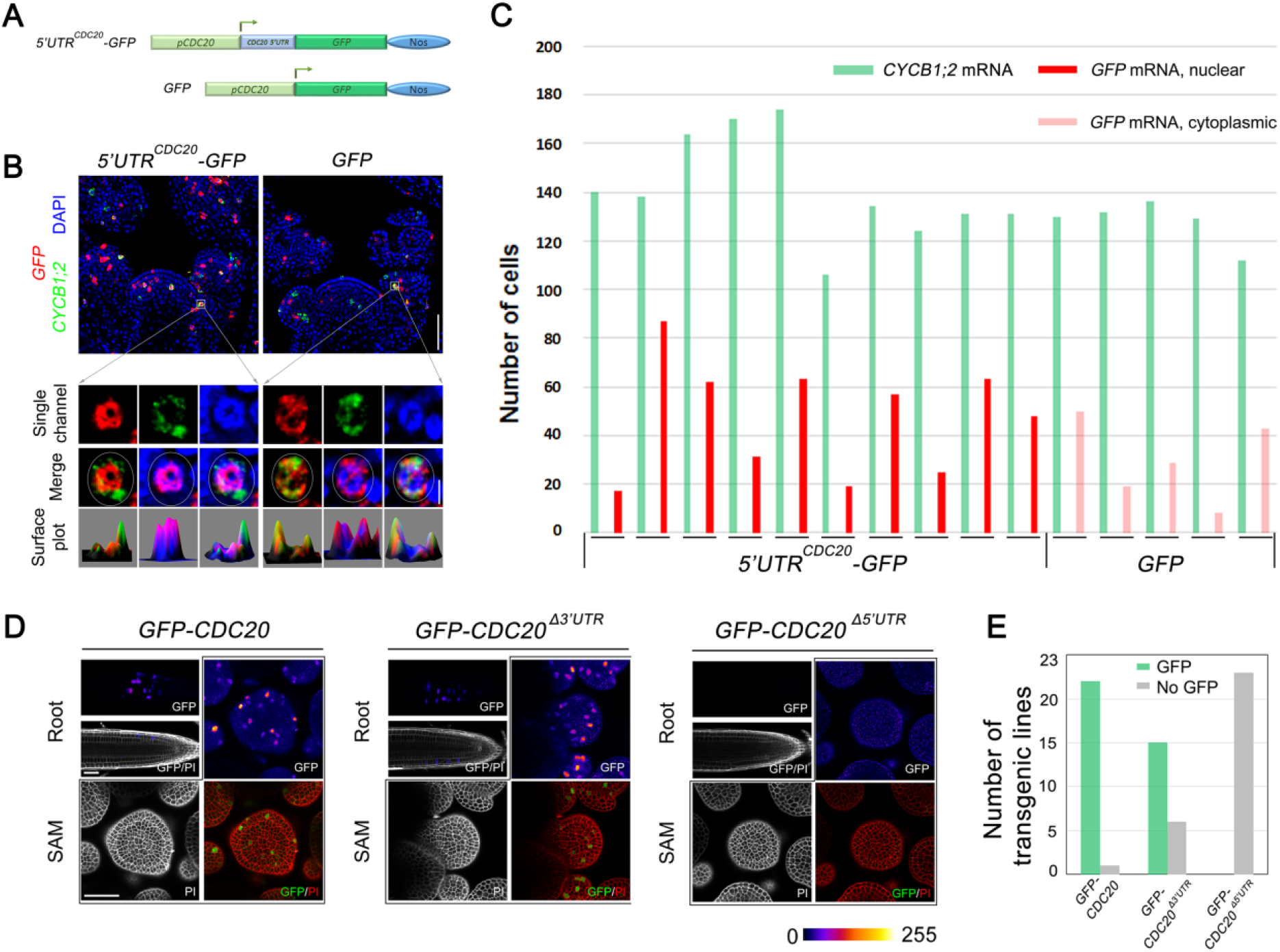
Dual Roles of 5’UTR in *CDC20* mRNA Nuclear Localization and Translation. (A) Schematic diagram of chimeric mRNA construction in which *GFP* was fused with *CDC20* 5’UTR. *GFP* alone was used a control. (B) Localization of *5’UTR^CDC20^-GFP* and *GFP* mRNAs in prophase cells. Scale bars, 50 µm for SAM overview (top panels) and 5 µm for single cells (bottom panels). (C) Quantification of the number of prophase cells expressing*5’UTR^CDC20^-GFP* and *GFP* mRNAs. Each pair of columns represents cell numbers from one meristem. (D) The expression of GFP-CDC20 fusion protein in root and SAM as revealed. No GFP fluorescence could be observed in 5’UTR truncated *GFP-CDC20* transgenic plants. Scale bar, 50 µm. (E) The number of transgenic lines analysed. GFP-CDC20 expression was detected in 22/23 lines of full length *GFP-CDC20* plants, 15/21 lines of 3’UTR truncated *GFP-CDC20* transgenic plants, and 0/23 of 5’UTR truncated *GFP-CDC20* transgenic plants.

### *CDC20* 5’UTR Is Required for Protein Translation

The cytoplasmic localization of *GFP-CDC20^Δ5’UTR^* mRNA in prophase cells, if translated, would be expected to interfere with proper cell cycle progression. However, we did not observe any cellular defect in chromosome alignment or segregation, and the transgenic plants grew normally. Confocal microscopy analysis revealed that in *GFP-CDC20^Δ3’UTR^* meristems, the fusion protein could be normally translated, showing clear GFP fluorescence in root and shoot apical meristem similar to the full length transcript (Figures 6D and 6E). However, no fluorescence could be observed in multiple independent *GFP-CDC20^Δ5’UTR^* transgenic lines, indicating that 5’UTR truncated *GFP-CDC20* mRNA cannot be properly translated. Taken together, these results demonstrate that the 5’UTR of *CDC20* plays dual roles in mRNA nuclear localization and translation.

## Discussion

To ensure the fidelity of chromosome segregation, APC/C activity needs to be precisely modulated, especially at early mitosis when APC/C targets (e.g. CYCB proteins) are playing crucial roles. Emi1 has been implicated in animals as the inhibitor of APC/C by binding to CDC20, preventing its interaction with APC/C substrates at prophase (Reimann et al., 2001). However, the role of Emi1 remains contentious as it was also shown to have little effect on APC/C^CDC20^ activity, and expression of a non-degradable version of Emi1 does not affect the destruction of cyclin A, cyclin B1 and securin (Di Fiore and Pines, 2007). Phosphorylation of APC/C subunits can facilitate CDC20 binding thus promoting APC/C activation (Sivakumar and Gorbsky, 2015). In mammalian cells APC/C phosphorylation is already initiated and CDC20 protein is also highly expressed at prophase (Kraft et al., 2003; Nilsson et al., 2008), which would presumably lead to APC/C activation. Therefore, it still remains obscure how APC/C activity is restrained during prophase. In plants, no Emi1 orthologue has been identified. GIG1/OSD1 and UVI4 have been suggested as the negative regulators of plant APC/C (Heyman et al., 2011; Iwata et al., 2011), but their direct effect on APC/C activity has not been determined. We found that in *Arabidopsis* dividing cells the mRNAs of *CDC20* and *CCS52B* are sequestered inside the nucleus. Nuclear retention of mRNAs is expected to block their accessibility to cytoplasmic ribosomes. Consistent with this scenario, neither CDC20 nor CCS52B proteins could be detected in prophase cells. As CDC20 and CCS52B are key activators of APC/C, it seems that absence of CDC20 and CCS52B proteins at prophase due to RNA nuclear sequestration would result in very low APC/C activity, thereby allowing cyclin B function (Figure S9).

Cellular mRNA localization has been proposed as a common mechanism to control local protein abundance. A systematic study revealed that 71% of expressed mRNAs in *Drosophila* embryos exhibit distinct cytoplasmic distribution patterns (Lécuyer et al., 2007). Compared to the predominant distribution in the cytoplasm, nuclear localization of protein coding mRNAs has rarely been encountered. Our data demonstrate that properly processed, unedited mature mRNAs can be specifically sequestered inside the nucleus, correlating with control (absence) of protein synthesis. Nuclear sequestration of *CDC20* and *CCS52B* mRNA, despite their high levels, prevents protein translation, but on the other hand could also generate a store of RNA molecules that can be rapidly released to the cytoplasm upon NEBD for protein synthesis, thus to efficiently activate APC/C.

RNA localization is guided by specific cis-acting elements that are mostly identified in the 3’UTR (Martin and Ephrussi, 2009). The localization signals contributing to the spatial distribution of *bicoid, nanos, xcat2*, β-actin mRNAs, and *histone* mRNAs have all been mapped to the 3’UTR (Iampietro et al., 2014; Martin and Ephrussi, 2009). However, deletion analysis revealed that the 3’UTR has no effect on *CDC20* mRNA nuclear localization. By contrast, when the 5’UTR is removed, the resulting *GFP-CDC20^Δ5’UTR^* chimeric mRNA is found to distribute into the cytoplasm. Furthermore, adding the *CDC20* 5’UTR was sufficient to sequester *GFP* mRNA in the nucleus, indicating that the 5’UTR is necessary and sufficient for *CDC20* mRNA nuclear sequestration. Despite being exported into the cytoplasm, the 5’UTR truncated *GFP-CDC20* RNA was not detectably translated, which is consistent with the important functions of 5’UTR in ribosome recruitment and translational initiation (Hinnebusch et al., 2016). Therefore, the dual roles of the 5’UTR in *CDC20* mRNA nuclear localization and translation provides a ‘belt-and-braces’ approach to avoid CDC20 protein synthesis and APC/C activation. RNA localization elements are recognized by trans-acting proteins. The RNA interactome capture method has been recently developed to identify *Xist* lncRNA binding proteins in human cells (Chu et al., 2015; McHugh et al., 2015; Minajigi et al., 2015). Applying this technology in plants to characterize *CDC20* and *CCS52B* mRNA interacting protein(s) would provide more insights into the understanding of mRNA localization, translational control, and cell cycle regulation.

## Materials and Methods

### Plant material and growth conditions

*Arabidopsis* Columbia ecotype (Col-0) was used as wild-type for the *in situ* hybridization analysis. The reporter lines GFP-MBD, H2B-RFP, CYCB1;1-GFP, and SUN2-GFP were described previously (Federici et al., 2012; Hamant et al., 2008; Oda and Fukuda, 2011; Reddy et al., 2005). Seeds were germinated on Murashige and Skoog agar plates and 7 day-old seedlings were transferred to soil. Plants were grown under long day conditions (16 h/8 h light/dark period) at 20 °C.

## mRNA *In Situ* Hybridization

### RNA Probe Synthesis

The cDNA fragments corresponding to each cell cycle gene were amplified with gene-specific primers (Table S4), ligated into the pGEM^®^-T Easy vector (Promega) and verified by sequencing. The plasmids containing the cDNA fragments were then used as templates for PCR with primers T7 and SP6. The PCR products were used as templates for in vitro transcription using the DIG RNA Labeling Kit (Roche). For fluorescence in situ probes, Fluorescein-12-UTP (Roche) was used instead of Digoxigenin-11-UTP (Roche).

### Sample Preparation

Shoot apices of *Arabidopsis* were harvested and fixed in FAA (3.7% formaldehyde, 5% acetic acid, 50% ethanol). The samples were embedded in wax and cut into 8-μm sections. The sections were processed by dewaxing, rehydration and dehydration, as described in (http://www.its.caltech.edu/~plantlab/protocols/insitu.pdf).

### Hybridization

The sections were hybridized with gene-specific probes at 55 °C. After washing with SSC, the slices were incubated with anti-digoxigenin-AP antibody (Roche) for 2 hours at room temperature. The signals were detected by overnight colour reaction at 28 °C using NBT/BCIP (Roche). Sense-strand hybridizations, yielding no hybridization with target mRNA, are shown as controls. Images were taken using a Zeiss AxioImager M2 microscope fitted with a Zeiss Axiocam MRc colour camera and a PlanApochromat 20x/ 0.8 NA objective.

### RNA Fluorescence *in situ* Hybridization (RNA FISH)

Samples were processed as above for *in situ* hybridization, except that anti-fluorescein-POD (Roche) or anti-digoxigenin-POD (Roche) antibodies were used. After antibody incubation, the hybridization signals were detected using TSA Plus Fluorescein Fluorescence System (Perkin Elmer) for green signals or TSA Plus Cy5 Fluorescence System (Perkin Elmer) for red signals. DAPI staining was performed by mounting the slices with 1µg/ml DAPI shortly before observing the *in situ* hybridization signals. Images were taken with a Zeiss LSM700 confocal microscope equipped with a 20 × 0.8NA dry objective. Laser excitation was 405 nm (DAPI), 488 nm (Fluorescein) and 633 nm (Cy5).

### Double RNA FISH

Double RNA FISH was used to check the mRNA expression of two genes in the same cells. Processed sections were hybridized with a mixture of two gene-specific probes, one labelled with digoxigenin and the other with fluorescein. The slices were incubated with anti-fluorescein-POD (Roche) and subsequently detected with TSA Plus Fluorescein Fluorescence System, giving green signals. After the first TSA reaction, 3% H_2_O_2_ (Sigma) was applied to quench peroxidase activity (1 hour incubation in 3% H_2_O_2_ was found to sufficiently quench all peroxidase activity of the first antibody). The slices were further incubated with anti-digoxigenin-POD antibody and detected by TSA Plus Cy5 Fluorescence System (Perkin Elmer), resulting in red signals.

### RNA FISH and Immunohistochemistry

RNA FISH was carried out as described above. After TSA-Cy5 reaction to reveal the mRNA hybridization signals, the sections were washed in PBST (PBS containing 0.3% v/v Triton X-100), and then blocked in PBS-Blocking buffer (PBS containing 1.0% bovine serum albumin, 0.2% powdered skim milk, and 0.3% Triton X-100) for 30 min at room temperature. The sections were then incubated with Alexa Fluor^®^ 488 conjugated GFP antibody (1:100 dilution) (A-21311, Molecular Probes) overnight at 4 °C. The slides were washed in PBST for 3 times, 5 min each and observed using a Zeiss LSM700 confocal microscope.

## Plasmid Construction and Plant Transformation

### *GFP* Fusion with Full Length *CDC20* and *CCS52B* mRNA

The MultiSite Gateway^®^ Three-Fragment Vector Construction system (Invitrogen) was used to generate plasmid constructs. For pCDC20.1::GFP-CDC20.1, a 2,417 bp promoter upstream of CDC20.1 ATG was amplified using genomic DNA as template with primers CDC20_promoter_F and CDC20_promoter_R. The PCR product was inserted into pDONR^TM^ P4-P1R by BP reaction, resulting in 1R4-pCDC20. The enhanced GFP (EGFP) coding sequence was amplified using primers GFP_gateway_F and GFP_gateway_R, and the product was inserted into pDONR^TM^ 221 by BP reaction, resulting in 221-GFP. A 3,161bp genomic fragment containing the whole coding sequence of CDC20.1 as well as 1,115bp 3’ region was amplified with primers CDC20_DNA_F and CDC20_DNA_R, and the PCR product was inserted into pDONR^TM^ P2R-P3, resulting in 2R3-gCDC20. The three entry constructs was incorporated into the binary vector pB7m34-GW by LR reaction. Similar strategy was applied to CCS52B. The primers used for CCS52B promoter were CCS52B_promoter_F and CCS52B_promoter_R; for coding region as well as 3’ region were CCS52B_DNA_F and CCS52B_DNA_R, and the constructs were named as 1R4-pCCS52B and 2R3-gCCS52B, respectively. pCDC20.1::GFP-CDC20.1 and pCCS52B::GFP-CCS52B were transformed into Arabidopsis wild-type Col-0 as well as nuclear reporter line H2B-RFP (Col-0 background) via Agrobacterium mediated transformation.

To construct *CDC20* cDNA fused with *GFP*, the full length cDNA including 5’ and 3’ UTR was first amplified from meristem cDNA library using primers CDC20_cDNA_F and CDC20_cDNA_R. *GFP* was amplified with primers GFP_F and GFP_R. *CDC20* cDNA and *GFP* fragments were ligated into pBluescript SK(-), resulting in SK-GFP-cCDC20, which was then incorporated into pB7m34-GW with *CDC20* promoter and Nos terminator by LR reaction.

### ***CDC20* 5’ UTR and 3’ UTR Deletions**

For 5’UTR deletion analysis, *CDC20* promoter was amplified with primers CDC20_promoter_F and CDC20_promoter_NoUTR_R. The PCR products were inserted into pDONR^TM^ P4-P1R by BP reaction, resulting in 1R4-pCDC20_No5’UTR. 1R4-pCDC20_No5’UTR was further introduced into the binary vector pB7m34-GW with 221-GFP and 2R3-gCDC20 by LR reaction. For 3’UTR deletion analysis, *CDC20* genomic sequence without 3’UTR was amplified with primers CDC20_KpnI and CDC20_SalI. *CDC20* terminator was amplified with primers CDC20_SalI_1 and CDC20_BamHI. The two fragments were ligated into pBluescript SK(-), and the resulting plasmid was used as template for PCR with primers CDC20_DNA_F and CDC20_DNA_R. The PCR product was inserted into pDONR^TM^ P2RP3, resulting in 2R3-gCDC20_NoUTR. 1R4-pCDC20, 221-GFP and 2R3-gCDC20_NoUTR were ligated into pB7m34-GW by LR reaction.

### *CDC20* Coding Sequence Deletions

Fusion PCR was used to generate *CDC20* ORF deletion constructs. Two PCR fragments with 25 bp overlapping were amplified with specific primers (Table S2). The PCR products were mixed and used as templates for a second round of PCR using primers GFP_GW1_F and GFPCDC20_GW1_R. The product was inserted into pDONR^TM^ 221 by BP reaction, and further incorporated into pB7m34-GW with *CDC20* promoter and Nos terminator by LR reaction.

### Observation of Fluorescent Reporter Expression by Confocal Microscopy

Shortly after bolting (stem length ~ 1 cm), the shoot apex was dissected and the fully developed flowers were carefully removed in order to expose the SAM. The meristem was then transferred to a square box containing fresh MS medium (Duchefa Biochemie - MS basal salt mixture) supplemented with vitamins (myoinositol 100 µg/ml, nicotinic acid 1 µg/ml, pyridoxine hydrochloride 1 µg/ml, thiamine hydrochloride 1 µg/ml, glycine 2 µg/ml) and 1% sucrose in 15 order to keep the meristem alive during observation. Viewed-stacks of SAMs were acquired with either a Zeiss LSM700 with 20 × NA 1.0 water dipping objective or a Leica SP8 with 25 × NA 1.0 water dipping objective. 3D rendering was carried out using either Zen (Zeiss) or LAS X (Leica) confocal microscope software. The cell boundaries of the SAM were revealed by 0.1% propidium iodide (PI) staining for 5 min. Laser excitations were 488 nm (PI, GFP) and 555nm or 561nm (RFP). GFP fluorescence intensity was measured in Fiji ImageJ. To display the fluorescence intensity as shown in Figures 5 and S7, the fluorescence pictures were edited with the LUT editor plugin in Fiji ImageJ.

For MG132 treatment, dissected meristems were emerged in liquid MS medium containing DMSO (Mock) or 50 µM MG132 (C2211 Sigma) for 2 hours. For time lapse experiment, dissected meristems were kept in MS medium (Duchefa) supplemented with vitamins and sucrose. The meristems were kept in growth chamber under long day conditions (16 h/8 h light/dark period) at 20 °C, and were taken out for confocal imaging at each time point.

## Acknowledgments

We would like to thank Jonathon Pines (The Institute of Cancer Research, London), David Ron (Cambridge Institute for Medical Research, University of Cambridge), Yrjö Helariutta and Henrik Jönsson (The Sainsbury Laboratory at Cambridge University), and Olivier Hamant (Plant Reproduction and Development Laboratory, INRA, ENS Lyon) for advice and insightful discussions. We also thank David E. Evans (Oxford Brookes University), Susan Armstrong (University of Birmingham), and Xinnian Dong (Duke University) for sharing seeds. We are grateful to Christoph Schuster for support with *in situ* hybridisation, Benoit Landrein for suggestions for confocal microscope analysis, Pawel Roszak for help with root sectioning, Alexis Peaucelle, Charles Melnyk, Paul Tarr, Pau Formosa Jordan and all members of the Meyerowitz Lab at the California Institute of Technology for helpful conversations. We appreciate Barbara Di Fiore and Anja Hagting (The Gurdon Institute, University of Cambridge), and Lisa Willis (The Sainsbury Laboratory at Cambridge University) for comments and careful reading of the manuscript. This work was funded by the Gatsby Charitable Trust (through fellowship GAT3395/DAA). E.M.M. is supported by the Howard Hughes Medical Institute and the Gordon and Betty Moore Foundation (through grant GBMF3406). R.W. and W.Y. are supported by the Leverhulme Trust (grant RPG-2015-285).

## Supplemental Figures

**Figure S1.**
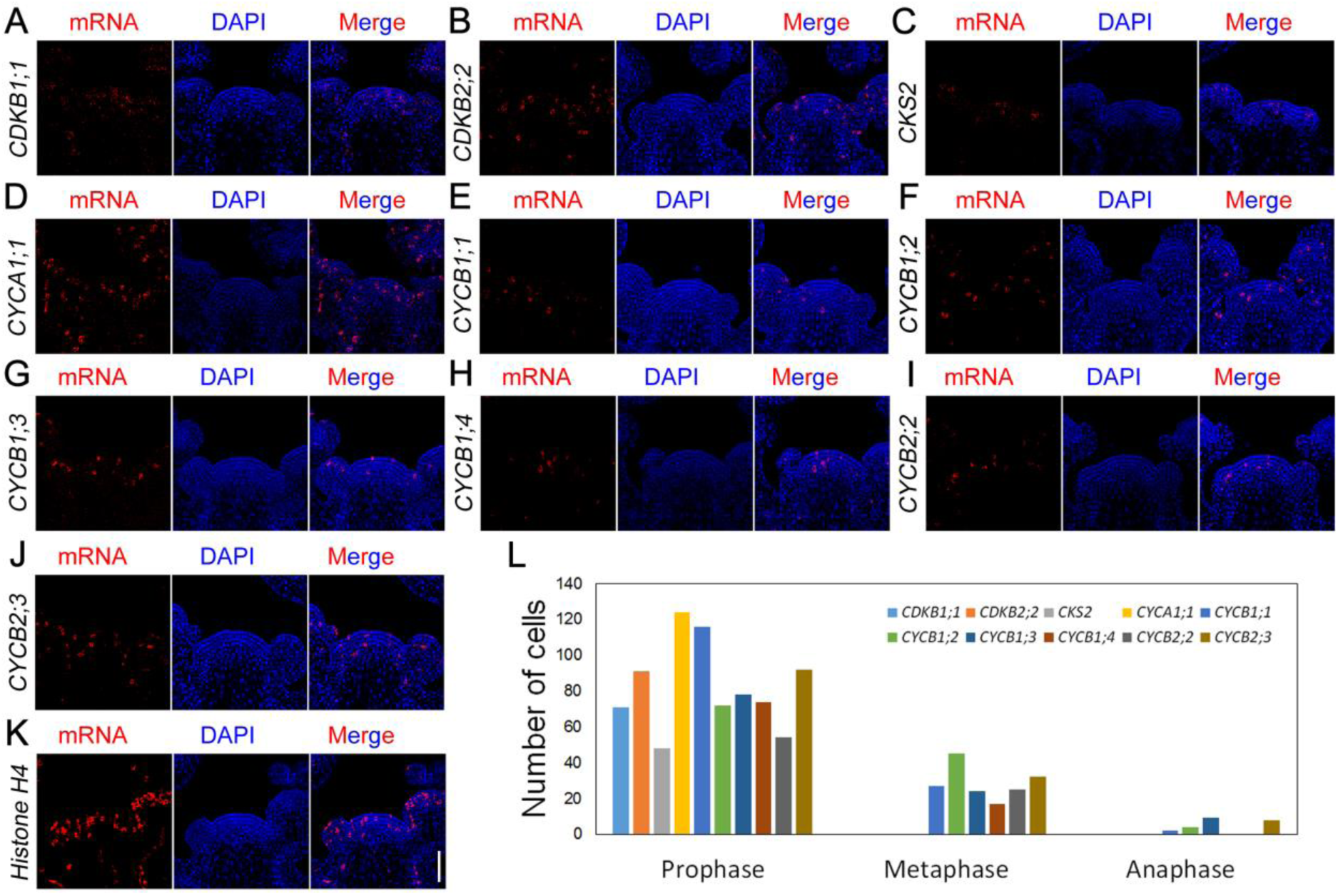
Mitosis Specific Expression of Cell Cycle Genes in the SAM. The mRNAs were detected by DIG labelled probes, which were further recognized by POD-labelled anti-DIG antibody coupled with the TSA-CY5 detection system. The nucleus was stained with DAPI. (A-K) Expression patterns of G2/M cell cycle genes. Scale bar, 50 µm. (L) Quantification of the number of cells expressing cell cycle genes at different mitotic stages.

**Figure S2.**
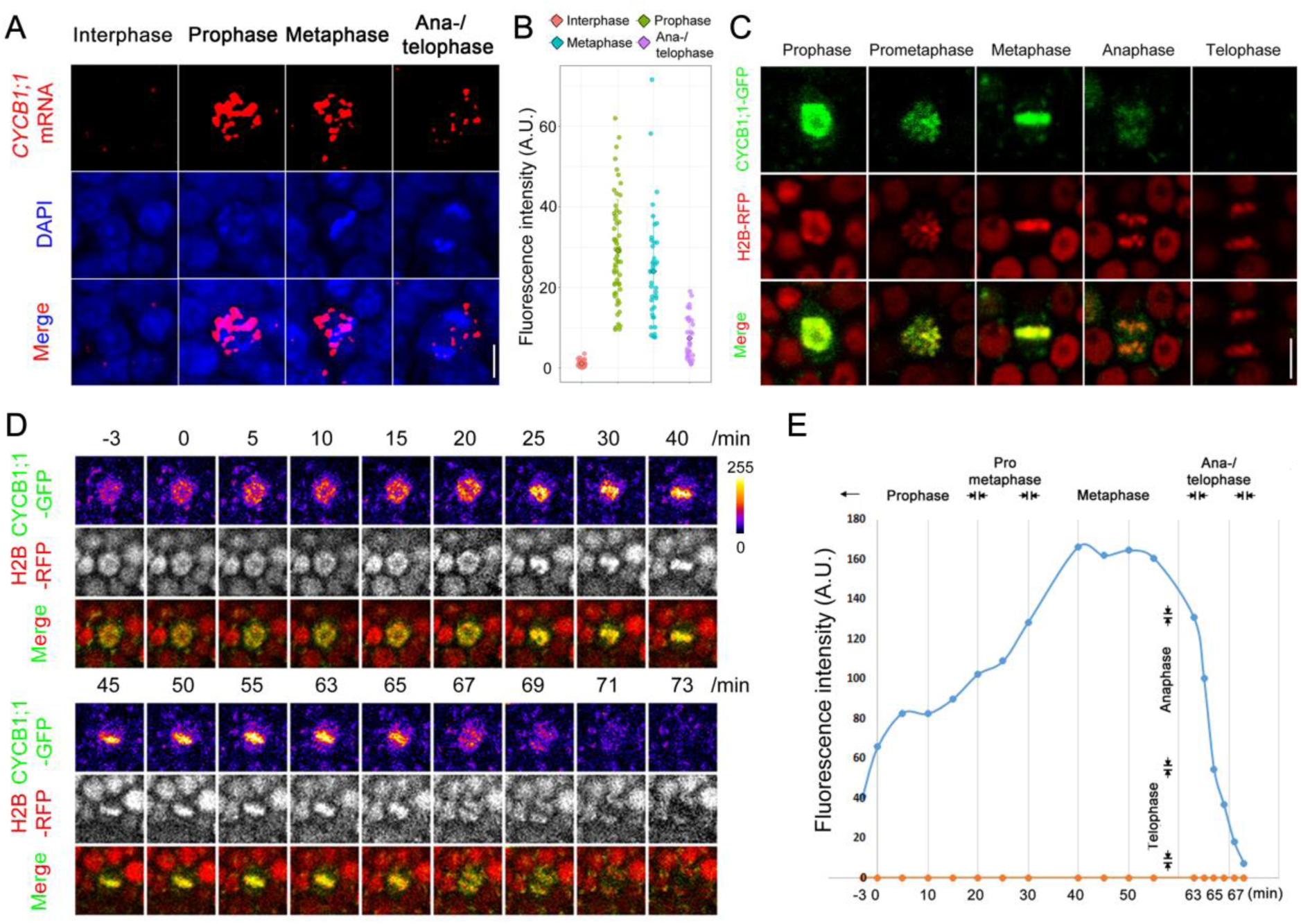
Rapid CYCB1;1 Protein Degradation at Metaphase-to-Anaphase Transition. (A) RNA FISH to show the accumulation of *CYCB1;1* transcripts at different stages of mitosis. Scale bar, 5 µm. (B) *CYCB1;1* mRNA levels at different stages of mitosis, as calculated from the fluorescence intensity of RNA FISH images. (C) CYCB1;1-GFP protein expression at different stages of the cell cycle. H2B-RFP is used to monitor chromosome alignment and segregation. Scale bar, 5 µm. (D and E) Protein dynamics of CYCB1;1-GFP during mitosis. GFP fluorescence intensity is shown in (E).

**Figure S3.**
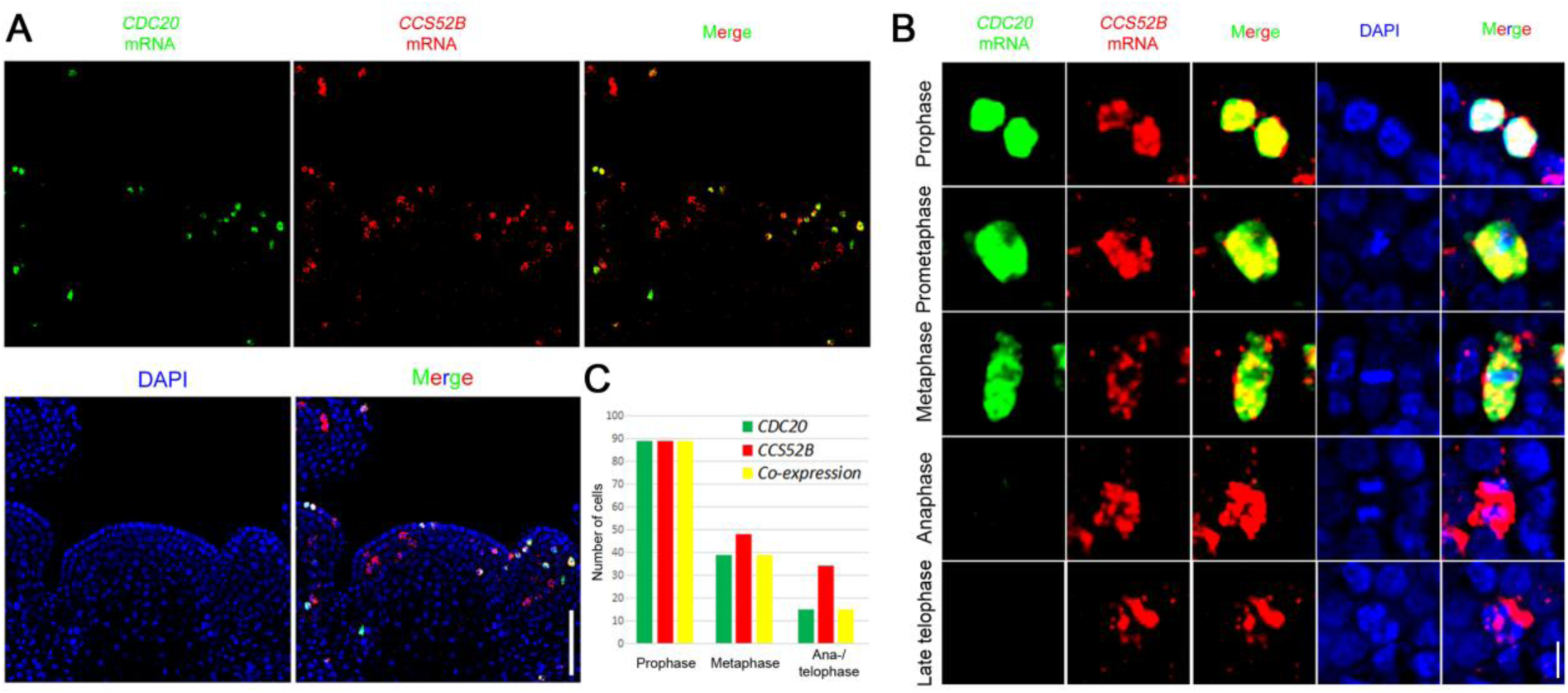
Co-expression Analysis of *CDC20* and *CCS52B*. (A) Double RNA FISH to show the expression patterns of *CDC20* and *CCS52B* in the same meristem. Scale bar, 50 µm. (B) Co-expression of *CDC20* and *CCS52B* at different mitotic stages. The anaphase and late telophase cells shown are those only expressing *CCS52B*. Scale bar, 5 µm. (C) Quantification of the number of cells that express *CDC20* and *CCS52B*.

**Figure S4.**
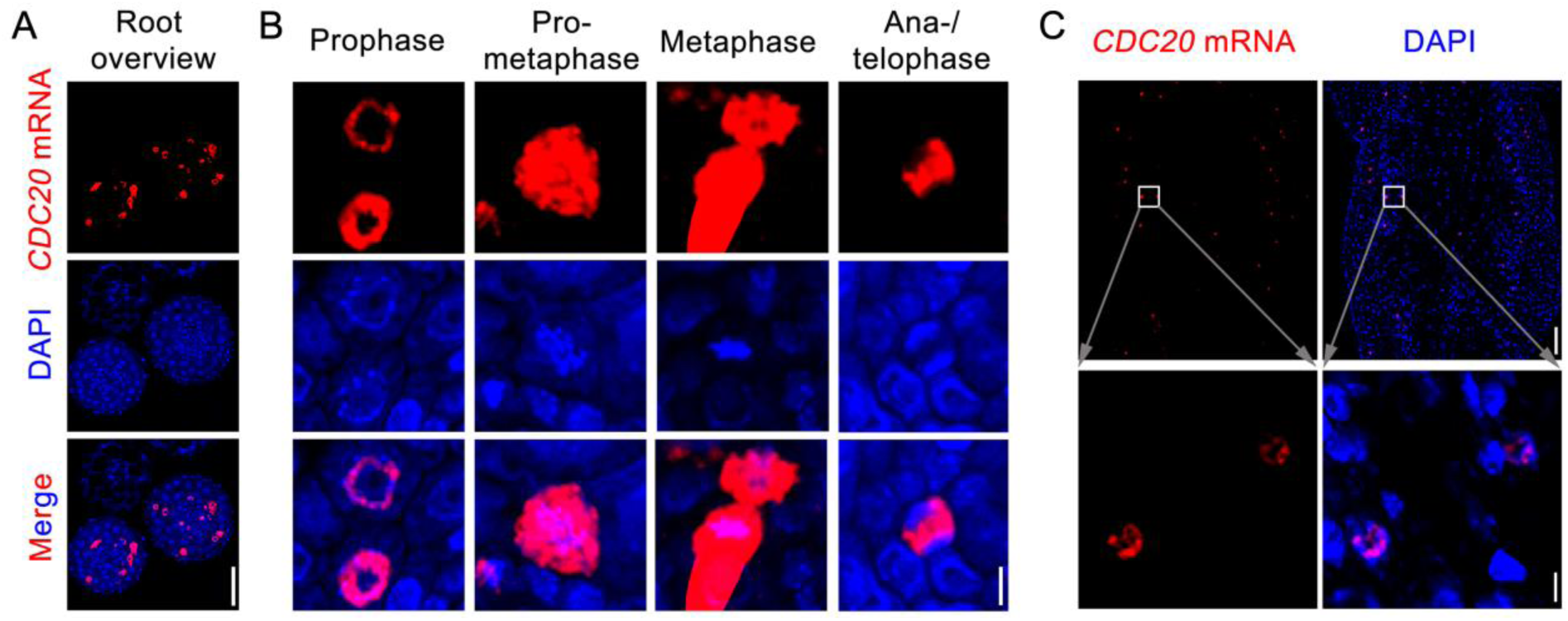
Expression Pattern of *CDC20* in Root and Shoot Dividing Cells. (A) Root overview. Scale bar, 50 µm. (B) Root cells at different stages of mitosis. Note that *CDC20* mRNA is sequestered inside the nucleus at prophase. Scale bar, 5 µm. (C) Nuclear localization of *CDC20* mRNA in shoot prophase cells. Scale bars, 50 µm for shoot overview (top panels) and 5 µm for cells (bottom panels).

**Figure S5.**
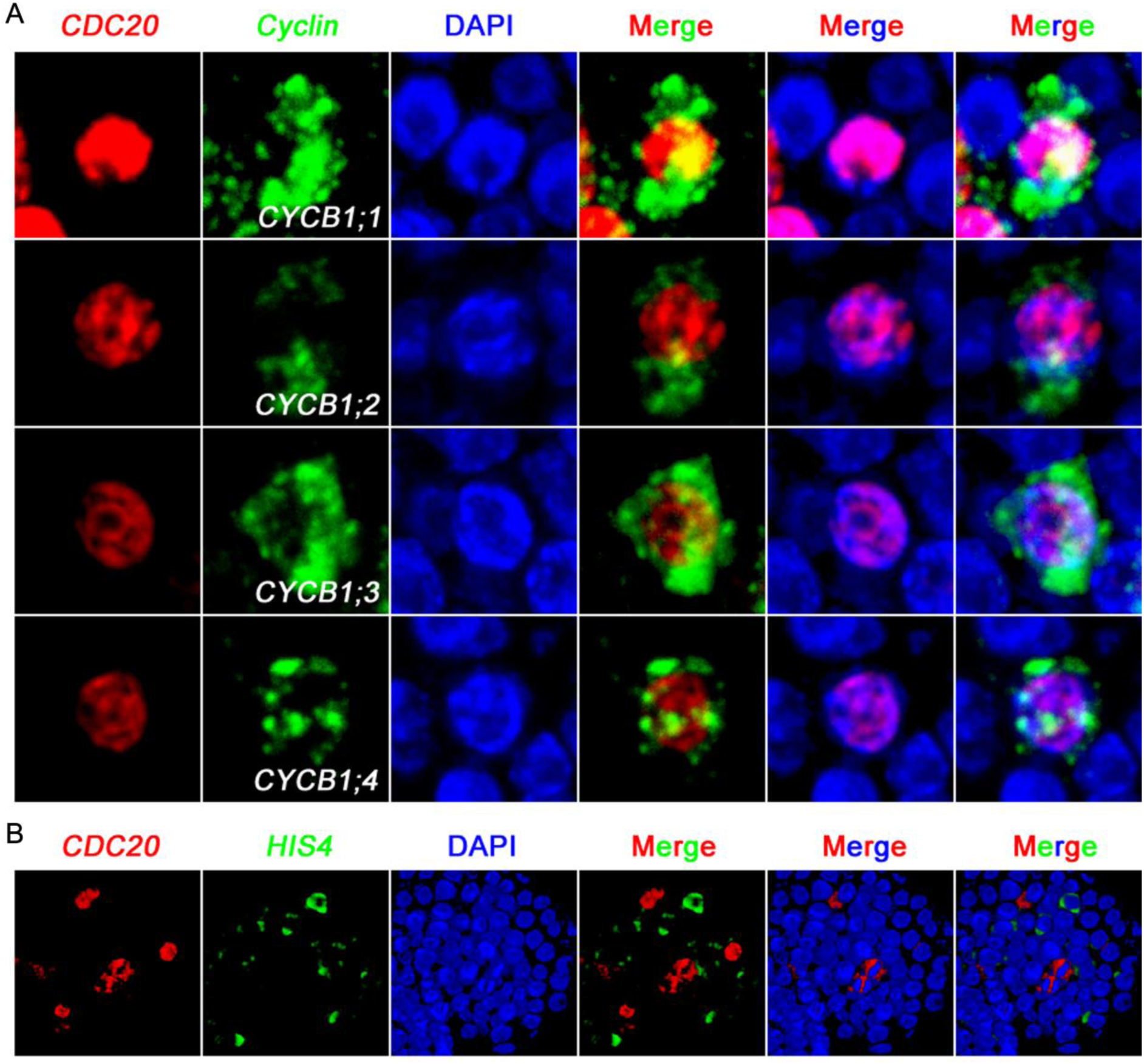
3-D Projection of Confocal Images to Show *CDC20* Expression Patterns with *CYCBs* and *HIS4* in the Same Meristems. (A) Nucleocytoplasmic separation of *CDC20* and *CYCB1* transcripts in prophase cells. (B) *CDC20* does not co-express with S-phase marker *HIS4* gene.

**Figure S6.**
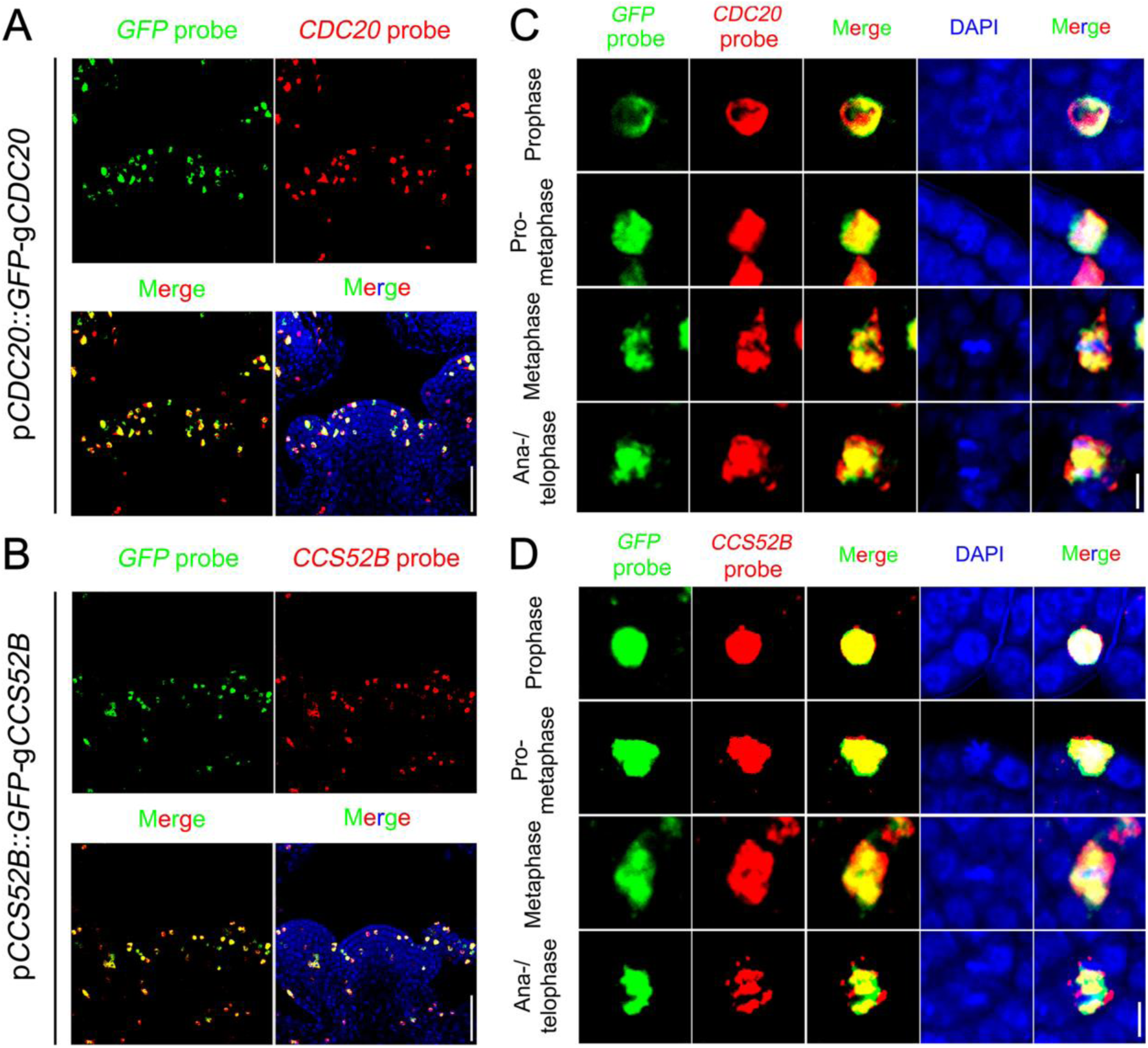
Fusion of *GFP* does not Affect *CDC20* or *CCS52B* mRNA Nuclear Localization. (A and B) Overview of *GFP* mRNA distribution with *CDC20* and *CCS52B* in *pCDC20::GFPCDC20* (A) and *pCCS52B::GFP-CCS52B* (B) transgenic plants. Scale bars, 50 µm. (C and D) Co-localization of *GFP* mRNA with *CDC20* (C) or *CCS52B* (D) mRNA in mitotic cells. Scale bars, 5 µm.

**Figure S7.**
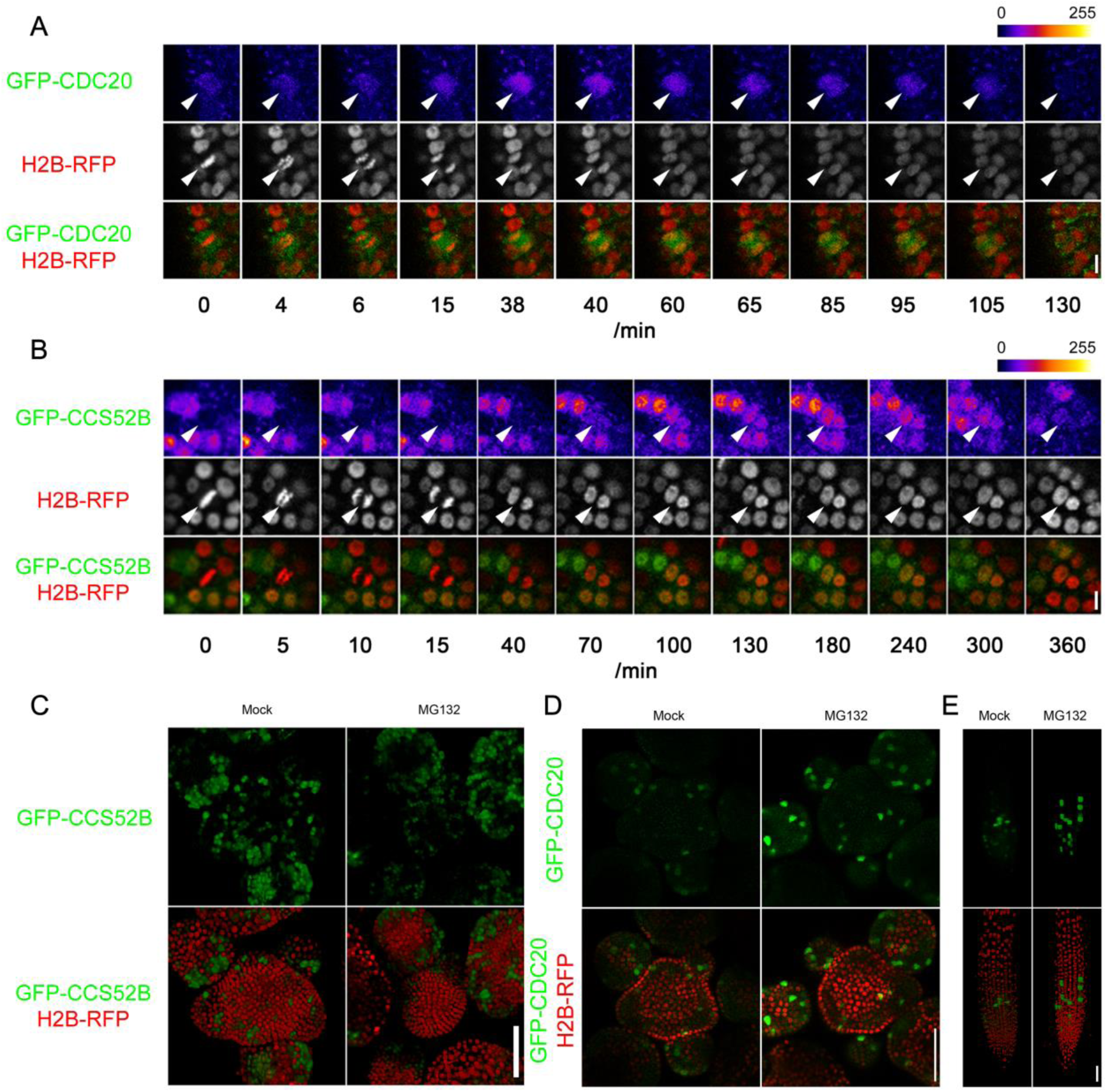
Fluctuation in the Protein Levels of CDC20 and CCS52B during the Cell Cycle. (A and B) Time-lapse imaging of GFP-CDC20 and GFP-CCS52B protein expression in the same cell as it undergoes division. Arrowheads indicate the cells analysed. Scale bars, 5 µm. (C) MG132 treatment does not affect GFP-CCS52B protein abundance. Scale bar, 50 µm. (D and E) The amount of GFP-CDC20 proteins in both SAM (C) and root (D) can be increased by MG132 treatment. Scale bar, 50 µm.

**Figure S8.**
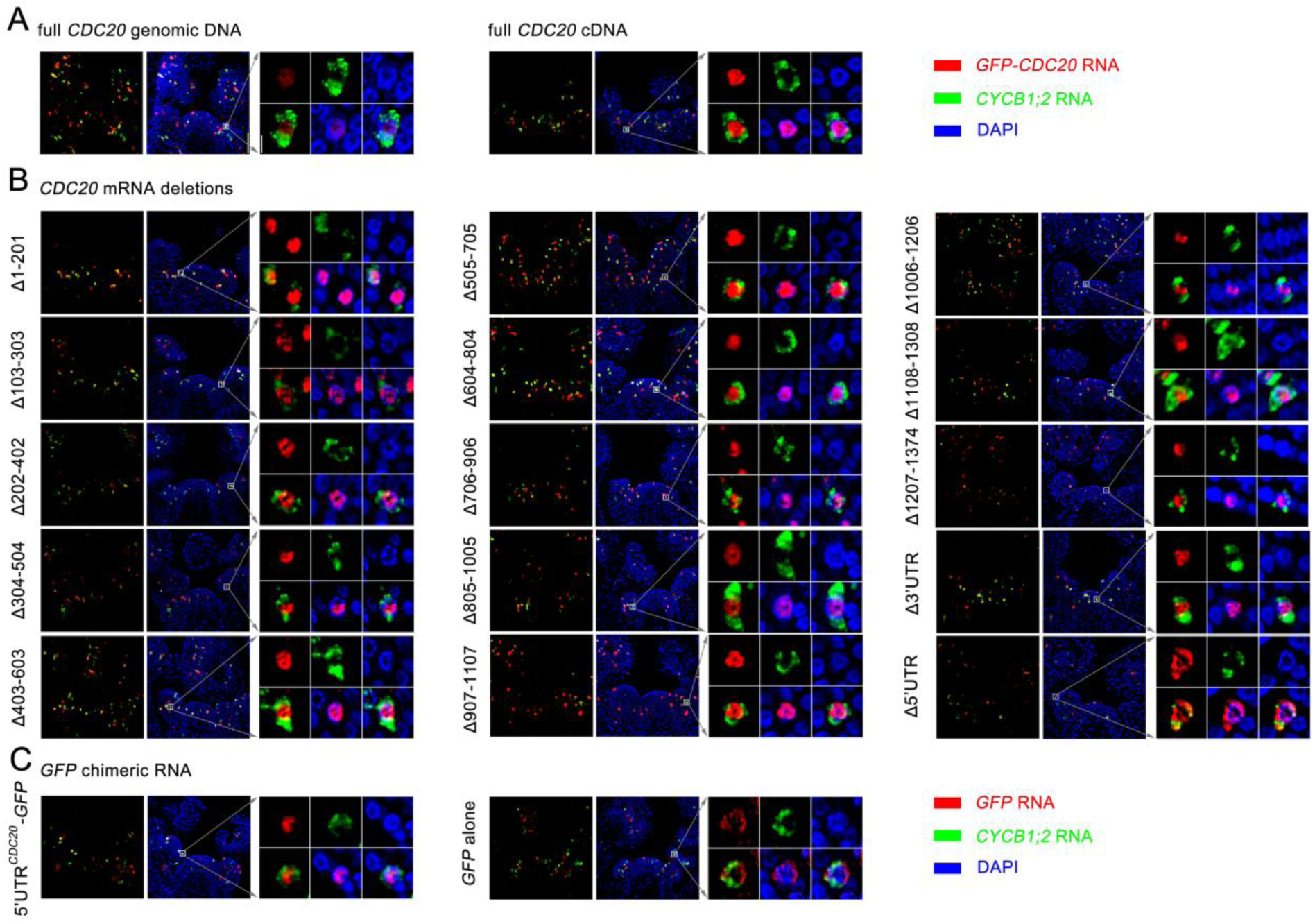
5’UTR Affects *CDC20* mRNA Nuclear Localization. (A) The expression patterns of full length *GFP-CDC20* mRNAs transcribed from genomic DNA or cDNA in the shoot apex. Shown are representative meristems from one of the independent transgenic lines. Scale bar, 50 µm for SAM overview and 5 µm for single cells. (B) The expression patterns of *GFP-CDC20* truncated mRNAs. (C) The expression patterns of *GFP* chimeric mRNAs.

**Figure S9.**
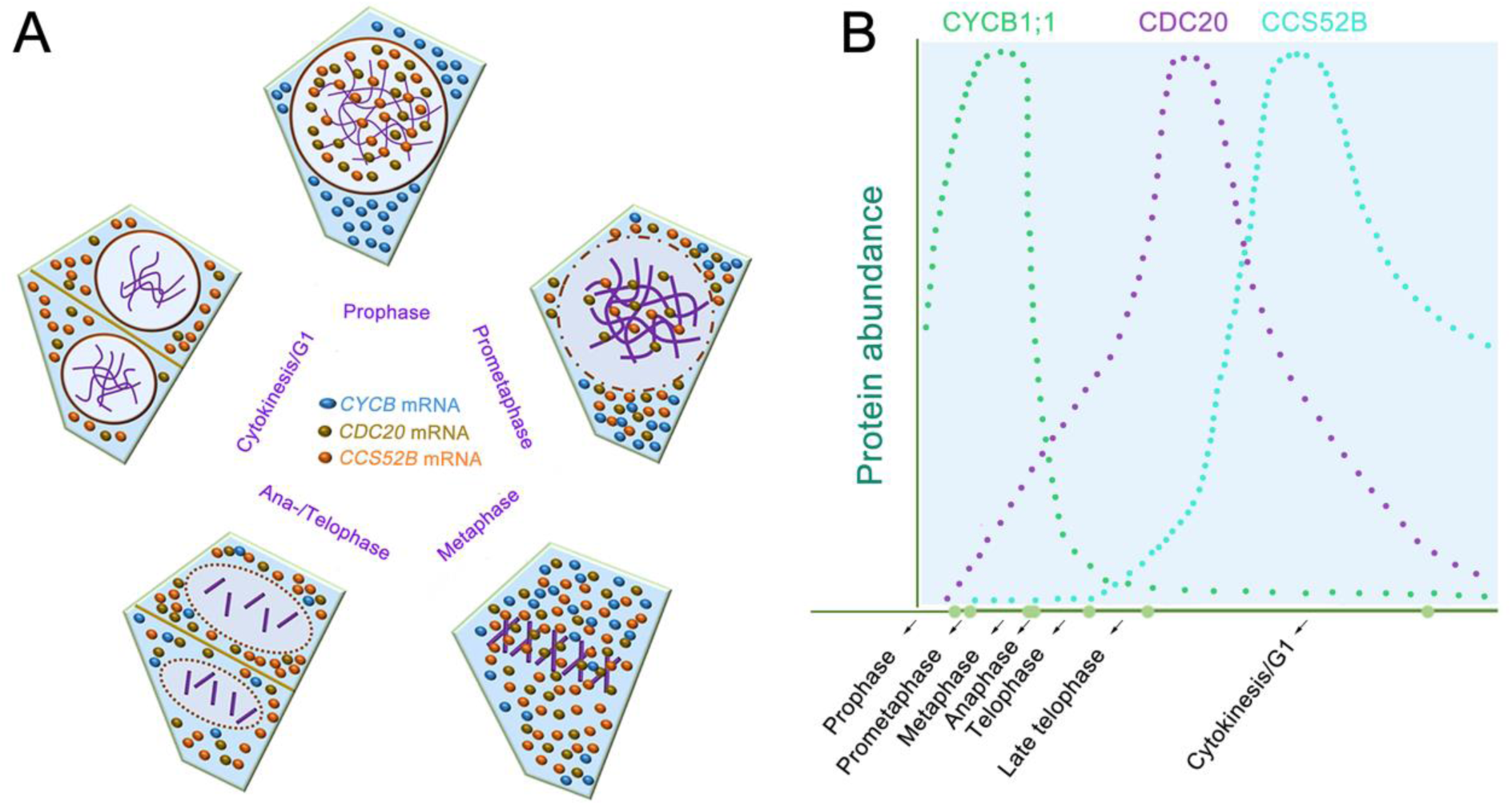
Model for Cell Cycle Control by mRNA Nuclear Sequestration. (A) Subcellular distribution of *CYCB*, *CDC20* and *CCS52B* mRNAs during cell cycle progression in plant stem cells. (B) CYCB, CDC20 and CCS52B protein dynamics. Nuclear sequestration of *CDC20* and *CCS52B* mRNAs in prophase prevents their translation to protein. Nuclear envelope breakdown at prometaphase enables redistribution of the mRNAs into the cytoplasm and subsequent protein synthesis, following which the proteins activate APC/C to destroy cyclin B proteins and other substrates.

